# CHARACTERIZATION OF EARLY-MATURING COWPEA (*Vigna unguiculata* (L.) WALP.) GERMPLASMS THROUGH ASSOCIATION ANALYSIS

**DOI:** 10.1101/2023.12.05.570049

**Authors:** Iduh Michael Ogbeche, Godspower Chibuike Ekeruo, T Vange, Lucky Omoigui, Odang Emmanuel Ogbu, Koletowo Faruq Oyinkansola, Ayeni Olalekan

## Abstract

Early maturing cowpea is a valuable crop for sub-Saharan Africa that offers a number of benefits, including a short growing season, drought tolerance, and the potential to improve food security. This study was conducted to analyze character associations in genetically diverse cowpea (*Vigna unguiculata* (L.) Walp.) germplasm, employing ten distinct accessions within a randomized complete block design with four replicates. Parameters such as days to first flowering, days to fifty percent flowering, number of branches per plant, days to ninety-five percent maturity, number of pods per plant, hundred-seed weight, total pod weight, and total seed weight were subjected to analysis of variance (ANOVA), Correlation, and Path Coefficient Analysis. Results revealed significant positive correlations between “branches per plant and pods per plant (r = 0.317, p ≤ 0,001),” “pods per plant and pod weight (r = 0.136, p < ≤ 0,001),” and “pod weight and seed weight (r =0.567, p ≤ 0,001).” Path coefficient analysis identified pod weight (0.840938) as the most influential direct contributor to yield, while the number of pods (-0.04268) indirectly influenced yield through pod weight. High grain yield selection can be achieved through indirect manipulation of pod weight. Accession IT99K-573-1-1, IT10K-863-11 and IT08K-125-24 exhibited the highest yield characters (pod weight and seed weight) and therefore further studies is recommended to consolidate this work, to enhance adoption by farmers.

## INTRODUCTION

Cowpea (*Vigna unguiculata* (L.) Walp.) 2n=22 is one of the principal food and cash crop legume grown in the semi-arid tropics covering Africa, Asia and Central America with great socio-economic, cultural, nutritional importance and a valuable component of the traditional cropping systems (FAO/WFP/IFAD 2022). Cowpea belongs to the family *Fabaceae* and sub-family *Faboideae* (Agbogidi, 2010). Other commonly used names of cowpea include southern pea, black eye pea, crowder pea, labia, niece, coupe, or frijole (Horn and Shimelis, 2020). Cowpea is a major food crop that is grown in over 100 countries and provides a source of income for over 40 million farmers. It is also a good source of protein, oil, vitamins, and minerals for millions of people, especially in developing countries (FAO 2022, NRIN, 2022). About 6.5 million metric tons of cowpea were reported produced annually on about 14.5 million hectares worldwide (Boukar *et al*., 2018). In Sub-Saharan Africa (SSA), West Africa is regarded as the major cowpea producing region with 80% of the total regional production reported for Nigeria and Niger at first and second positions respectively for 14 years in a row (Aboki, and Yuguda, 2013; Huynh *et al*., 2016; FAO 2022). In addition, Nigeria has been the leading producer and consumer of cowpea globally with an approximately 3.3 million tonnes (FAO 2022).

Cowpea production is generally low as a result of some factors such as lack of improved varieties, disease, drought, insect pest and weeds among other factors (Gungula and Garjila, 2005). The production level is also not enough as food production is lagging behind population growth; there is also soaring demand for livestock feeds and the deficiency of protein in the dietary requirement of average Nigerians. Therefore, it becomes imperative to increase to an exponential rate the yield of cowpea production in Nigeria (Obi *et al*., 2017; Ajisegiri *et al*., 2020)

Yield, being a complex trait, is influenced by various contributing factors controlled by polygenes and environmental conditions. Therefore, it is essential to gain insights into the associations among these factors, whether direct or indirect, to understand their impact on yield (Virk and Bansal 2000). While correlation coefficients elucidate the degree of association, interpreting such associations becomes challenging as indirect connections between factors increase (Virk and Bansal 2000; Piepho, and Ogata, 2022). In such cases, the path coefficient analysis, developed by Wright (1921), proves valuable. This method helps dissect the total correlation coefficient into direct and indirect effects, providing a clearer understanding of each component trait’s relative contribution to the final yield. Therefore, employing correlation and path analysis in the collection, maintenance, and evaluation of germplasm is a fundamental step in initiating a breeding program for the genetic enhancement of cowpea yield (Kashiwagi *et al*., 2012; Borner 2020). This research aimed at studying the correlation between yield and its attributing parameters and also to estimate direct and indirect effect of yield contributing characters on early-maturing cowpea seed yield.

## MATERIALS AND METHODS

The study was conducted during the 2020 cropping season at the College of Agronomy Teaching and Research Farm, Joseph Sarwuan Tarka University, Makurdi located at Latitude 7°14’ North and longitude 8°24’ East, elevation 98m, which falls within the Southern Guinea Savannah agro-ecological zone of Nigeria. Seeds of cowpea germplasms were obtained from the International Institute of Tropical Agriculture (IITA) through the Molecular Biology Laboratory, Joseph Sarwuan Tarka University, Makurdi. Lists of accessions are presented in Table 1. The experiment was laid out in a Randomized Complete Block Design (RCBD) with three (3) replications. The experimental area was ploughed and harrowed. Field was marked and demarcated into three replications spaced 1m apart. Each replication consists of ten plots and each plot contains four rows at 4m in length, spaced at 0.75m apart. Harvesting was done manually, when the pods have turned straw brown, that is 95% of the pods had matured for each variety.

**Table 1:** List of Early-Maturing Cowpea Germplasms Characterized Through Association Analysis.

| S/N | Germplasms | Source |
| --- | --- | --- |
| 1 | IT08K-180-10 | IITA |
| 2 | IT08K-125-24 | IITA |
| 3 | IT08K-150-27 | IITA |
| 4 | IT10K-822-7 | IITA |
| 5 | IT10K-822-9 | IITA |
| 6 | IT10K-863-11 | IITA |
| 7 | IT10K-961-7 | IITA |
| 8 | IT10K-817-7 | IITA |
| 9 | IT10K -817-3 | IITA |
| 10 | IT99K-573-1-1 | IITA |

Data was collected and recorded for analysis from each individual plot on the following parameters: days to first flowering, days to 50 % flowering, number of branches per plant, 95% pod maturity, numbers of pods per plant, 100 seed weight, total pod weight, and total seed weight. All data collected were subjected to analysis of variance (ANOVA) using the Genstat Statistical Software 17th edition. Means were separated using the turkey pairwise comparison at 95% level of significance, where significant means exist. Pearson’s correlation coefficient was used to test for a correlation between all parameters measured. Path coefficient analysis was performed using Microsoft Excel 2016.

## RESULTS

Mean squares from analysis of variance for yield and yield related traits in early-maturing cowpea germplasms is presented in Table.2. The result showed that significant variations were observed among the ten (10) cowpea germplasms for days to first flowering, days to fifty percent flowering, number of branches per plant, days to ninety-five percent maturity, hundred seed weight, and total seed weight. There was however no significant variation observed for number of pods per plant and total pods weight.

**Table 2:** Analysis of Variance for Yield and its Components in Early Maturing Cowpea Germplasms.

| Sources of variations | Degree of freedom | Days to first flowering | Days to 50% Flowering | Numbers of branches per plant | Days to 95% Maturity | Numbers. of pod per plant | 100 seed weight | Total pod weight | Total seed weight |
| --- | --- | --- | --- | --- | --- | --- | --- | --- | --- |
| Replication | 2 | 1.30 | 2.43 | 1.43 | 11.70 | 1235.70 | 0.34 | 143628.00 | 18951.00 |
| Variety | 9 | 14.73** | 24.46** | 3.19** | 49.12** | 623.80<sup>ns</sup> | 21.53** | 66116.00<sup>ns</sup> | 27798.00** |
| Error | 18 | 1.45 | 1.80 | 0.84 | 2.96 | 295.80 | 0.43 | 41416.00 | 4084.00 |
| Total | 29 |  |  |  |  |  |  |  |  |
| CV% |  | 3.20 | 3.10 | 21.20 | 2.60 | 21.40 | 3.90 | 18.40 | 8.40 |

**KEY:** \*\* = Highly significant at $P \leq 0.01$ , ns = not significant, CV = Coefficient of variation,

The mean performance for ten early-maturing cowpea genotypes are presented in Table 3. Days to first flowering ranged from 34.67-41.00 with an overall mean of 37.60 days (Table 3). Cowpea accession IT08K-125-24, IT08K-961-7, IT08K-863-11, IT99K-573-1-1 and IT08K-180-10 are among the earliest flowering germplasms within the range of 34-37 days after planting, while cowpea accession IT10K-817-3, and IT10K-817-7 took longer days to flower (≥40 days) as shown in Table.3. The variability observed for days to fifty percent flowering was moderate, as reflected by its means Table 3, ranging from 39.33 days after planting for IT08K-125-24 to 47.33 days after planting for IT10K-817-7. IT08K-125-24 and IT08K-863-11 attained fifty percent flowering < 40 days after planting while other germplasm took more than 40 days to attain fifty percent flowering. The variation for numbers of branches per plant among the cowpea germplasms was moderate as reflected by the mean Table 3. Cowpea accession IT10K-961-7 and IT08K-125-24 had the least numbers of branches 3.000 each while cowpea accession IT10K817-7 and IT08K150-27 had the highest numbers of braches per plant, 5.667 and 6.000 respectively. Significant variations in the number of days to ninety-five percent (95%) maturity is presented in table 3. The mean value ranged from 59.33 days after planting to 71.67 days after planting. Cowpea accession IT10K-863-11 matured first, after 59.33 days of planting, while cowpea accession IT10K-822-7 took more days to mature, 71.67 days after planting. The overall mean value for this variable is 67.10 days. The mean estimate for numbers of pods per plant among the ten early-maturing cowpea germplasms is also presented in table 3. The average numbers of pod ranged from 55.00 to 100.67. Cowpea accessions IT08K150-27, IT10K-822-7, IT99K-573-1-1, IT08K-180-10 and IT10K-817-3 all produced pods ≥ 85. The weight of hundred seeds among the early-maturing cowpea accessions ranged from 12.13g to 20.20g with Cowpea accessions IT08K150-27, IT10K-822-7, IT10K-822-9, and IT99K-573-1-1 weighing between 18g to 20.20g while cowpea accessions IT10K-863-11 and IT10K-961-7 weighed the least 12.13g and 14.70g respectively.

**Table 3:** Mean Performance for Yield and Yield Component of Early-Maturing Cowpea Germplasms.

| Germplasms | Days to first flowering | Days to 50% Flowering | Numbers of branches per plant | Days to 95% Maturity | Numbers. of pod per plant | 100 seed weight | Total pod weight | Total seed weight |
| --- | --- | --- | --- | --- | --- | --- | --- | --- |
| IT08K-180-10 | 36.33<sup>cd</sup> | 41.00<sup>de</sup> | 3.667<sup>bc</sup> | 67.67<sup>b</sup> | 87.00<sup>abc</sup> | 15.53<sup>d</sup> | 1212<sup>ab</sup> | 786.3<sup>b</sup> |
| IT08K-125-24 | 34.67<sup>d</sup> | 39.33<sup>e</sup> | 3.000<sup>c</sup> | 64.00<sup>c</sup> | 65.33<sup>bc</sup> | 15.13<sup>d</sup> | 1193<sup>ab</sup> | 812.8<sup>b</sup> |
| IT08K-150-27 | 39.33<sup>ab</sup> | 44.67<sup>b</sup> | 6.000<sup>a</sup> | 69.67<sup>ab</sup> | 100.67<sup>a</sup> | 19.77<sup>ab</sup> | 938<sup>b</sup> | 638.3<sup>c</sup> |
| IT10K-822-7 | 39.33<sup>ab</sup> | 45.00<sup>ab</sup> | 5.000<sup>ab</sup> | 71.67<sup>a</sup> | 96.33<sup>ab</sup> | 19.63<sup>ab</sup> | 1076<sup>ab</sup> | 644.8<sup>c</sup> |
| IT10K-822-9 | 38.00<sup>bc</sup> | 43.67<sup>bc</sup> | 4.667<sup>abc</sup> | 69.00<sup>ab</sup> | 71.00<sup>abc</sup> | 20.20<sup>a</sup> | 1001<sup>ab</sup> | 637.9<sup>c</sup> |
| IT10K-863-11 | 35.67<sup>d</sup> | 39.67<sup>de</sup> | 3.667<sup>bc</sup> | 59.33<sup>d</sup> | 75.33<sup>abc</sup> | 12.13<sup>e</sup> | 839<sup>b</sup> | 825.6<sup>ab</sup> |
| IT10K-961-7 | 35.33<sup>d</sup> | 40.00<sup>de</sup> | 3.000<sup>c</sup> | 61.67<sup>cd</sup> | 75.00<sup>abc</sup> | 14.70<sup>d</sup> | 1117<sup>ab</sup> | 773.0<sup>b</sup> |
| IT99K-573-1-1 | 36.33<sup>cd</sup> | 42.00<sup>cd</sup> | 4.333<sup>abc</sup> | 69.67<sup>ab</sup> | 91.67<sup>ab</sup> | 18.73<sup>bc</sup> | 1353<sup>a</sup> | 930.0<sup>a</sup> |
| IT10K-817-3 | 40.00<sup>ab</sup> | 46.00<sup>ab</sup> | 4.333<sup>abc</sup> | 68.33<sup>b</sup> | 85.00<sup>abc</sup> | 15.67<sup>d</sup> | 1192<sup>ab</sup> | 804.7<sup>b</sup> |
| IT10K-817-7 | 41.00<sup>a</sup> | 47.33<sup>a</sup> | 5.667<sup>a</sup> | 70.00<sup>ab</sup> | 55.00<sup>c</sup> | 17.67<sup>c</sup> | 1113<sup>ab</sup> | 719.4<sup>bc</sup> |
| GRAND MEAN | 37.60 | 42.87 | 4.33 | 67.10 | 80.20 | 16.92 | 1103.00 | 757.00 |
| LSD | 2.06 | 2.30 | 1.57 | 2.95 | 29.50 | 1.13 | 349.10 | 109.60 |

**Note:** Mean with same or similar alphabets are not statistically different from each other

The results also showed that total pod weight varied significantly among the early-maturing cowpea germplasms evaluated. Cowpea accession IT10K-863-11 and IT10K-150-27 weighed the least 839kh/ha-^1^ and 938kh/ha-^1^ respectively, while cowpea accession IT08K-125-24, IT08K-180-10 and IT99K-573-1-1 weighed the most 1193kg/ha-^1^, 1212kg/ha-^1^ and 1353kg/ha-^1^ respectively. The grand mean for this character is 1103.00kg/ha. The mean value for total seed weight ranged from 637.9kg/ha for cowpea accession IT10K-822-9 to 930.0kg/ha for cowpea accession IT99K-573-1-1. with an overall mean value of 757kh/ha-^1^.

Pearson’s correlation coefficient between the various characters is presented in Table 4. The result shows that total seed weight was negative and not significantly correlated with all parameters observed except total pod weight (r = 0.567, P ≤ 0.001), which was positive and significantly correlated with total seed weight. Total pod weight was positive and significantly correlated with all parameters except numbers of branches per plant (r = -0.097) which was negative and not significantly correlated. Among the traits, the highest positive correlation of trait with total pod weight were numbers of days to ninety-five percent maturity (r = 0.164, P ≤ 0.001) and numbers of pods per plant (r = 0.136, P ≤ 0.001), while days to first flowerings (r = 0.061, P ≤ 0.005), days to fifty percent flowering (r = 0.074, P ≤ 0.005) and hundred seed weight (r = 0.016, P ≤ 0.005) were significantly correlated.

**Table 4:** Pearson’s Correlation coefficients for agro-morphological traits evaluated in early maturing Cowpea Germplasms.

|  | Days to first flowering | Days to 50% Flowering | Numbers of branches per plant | Days to 95% Maturity | Numbers of pod per plant | 100 seed weight | Total pod weight | Total seed weight |
| --- | --- | --- | --- | --- | --- | --- | --- | --- |
| Days to first flowering | 1 |  |  |  |  |  |  |  |
| Days to 50% Flowering | 0.905** | 1 |  |  |  |  |  |  |
| Numbers of branches per plant | 0.646** | 0.751** | 1 |  |  |  |  |  |
| Days to 95% Maturity | 0.582** | 0.589** | 0.495** | 1 |  |  |  |  |
| Numbers. of pod per plant | -0.036<sup>ns</sup> | 0.083* | 0.317** | 0.111** | 1 |  |  |  |
| 100 seed weight | 0.465** | 0.536** | 0.592** | 0.785** | 0.226** | 1 |  |  |
| Total pod weight | 0.061* | 0.074* | -0.097<sup>ns</sup> | 0.164** | 0.136** | 0.016* | 1 |  |
| Total seed weight | -0.393<sup>ns</sup> | -0.403<sup>ns</sup> | -0.488<sup>ns</sup> | -0.318<sup>ns</sup> | 0.046<sup>ns</sup> | -0.499<sup>ns</sup> | 0.567** | 1 |

**KEY:** \*\* = Highly significant at $P \leq 0.01$ , \* = Significant at $P \leq 0.05$ , ns = not significant.

Hundred seed weight was positive and significantly correlated with days to first flowering (r = 0.465, P ≤ 0.001), days to fifty percent flowering (r = 0.536, P ≤0.01), numbers of branches per plant (r = 0.592, P ≤ 0.001) days to ninety-five percent maturity (r = 0.785, P ≤ 0.001) and numbers of pods per plant (r = 0.226, P ≤ 0.001). Number of pods per plant was positive and significantly correlated all traits except numbers of days to first flowering (r = -0.036), which was negative and not significantly correlated to numbers of pods per plant. Among the traits, the highest positive correlation of trait with numbers of pods per plants were, Days to ninety-five percent maturity (r = 0.111, P ≤ 0.001) and numbers of branches per plant (r = 0.317, P ≤ 0.001). Days to ninety-five percent maturity was positive and significantly correlated with days to first flowering (r = 0.582, P ≤ 0.001), days to fifty percent flowering (r = 0.589, P ≤ 0.001) and numbers of branches per plant (r = 0.495, P ≤ 0.001). Numbers of branches per plant was positive and significantly correlated with days to first flowering (r = 0.646, P ≤ 0.001) and days to fifty percent flowering (r = 0.751, P ≤ 0.001).

The direct and indirect effect of Seven dependent characters on seed yield per plant as an independent character was obtained in path coefficient analysis using genotypic and correlation coefficient are presented in Table 5. The highest positive direct effect on seed yield was exhibited by total pod weight (0.840938) followed by number of branches per plant (0.079081) and days to ninety-five percent maturity (0.054855) whereas days to first flowering (-0.14532), days to fifty percent flowering (- 0.06231), numbers of pods per plant (-0.04268) and hundred seed weight (-0.22416) were contributing negative direct effect on seed yield.

**Table 5:** Path Coefficients Showing Direct (bold) and Indirect (not bold) Effect on Yield in Early-Maturing Cowpea Germplasms.

|  | Days to first flowering | Days to 50% Flowering | Numbers of branches per plant | Days to 95% Maturity | Numbers of pod per plant | 100 seed weight | Total pod weight |
| --- | --- | --- | --- | --- | --- | --- | --- |
| Days to first flowering | **-0.14532** | -0.05644 | 0.05201 | 0.031905 | 0.001803 | -0.10914 | -0.187 |
| Days to 50% Flowering | -0.13162 | **-0.06231** | 0.058997 | 0.032333 | -0.00356 | -0.12022 | -0.17677 |
| Numbers of branches per plant | -0.09557 | -0.04648 | **0.079081** | 0.027154 | -0.01239 | -0.13423 | -0.33497 |
| Days to 95% Maturity | -0.08452 | -0.03673 | 0.039147 | **0.054855** | -0.00475 | -0.02493 | -0.10974 |
| Numbers. of pod per plant | -0.006137 | -0.0052 | 0.022966 | 0.006101 | **-0.04268** | -0.05067 | 0.108962 |
| 100 seed weight | -0.07075 | -0.03342 | 0.047356 | 0.043082 | -0.00965 | **-0.22416** | -0.25113 |
| Total pod weight | 0.032315 | 0.013098 | -0.0315 | -0.00716 | -0.00553 | 0.06694 | **0.840938** |

## DISCUSSION

Analysis of variance for all characters observed showed high significant variability among the varieties. This variation observed among the ten cowpea germplasms evaluated may be attributed to the different genetic makeup of the genotypes studied which gives an ample scope for improvement in population through various breeding approaches. Cowpea accession IT08K-125-24 was the first to flower and also the first to attain fifty percent maturity. Whereas cowpea accession IT10K-817-7 flowered last and was also the last to attain fifty percent flowering among the germplasms evaluated, these variations in flowering could be as a result of the inherent genetic traits embedded in the germplasms or could be environmental factors, this work agrees with Egbe *et al*., (2010 who observed similar trend among 56 cowpea germplasms and reported that flowering ranges between 30 – 45 days after planting, they further reported that this variability is due to genetic and environmental factors. Seed yield and yield related traits revealed high significant variations among all germplasms studied, Cowpea accession IT99K-573-1-1 weighed more for pod weight and seed weight than all other genotypes evaluated, while cowpea accession IT10K-822-9 weighed the least for total seed weight. This result is slightly in tandem with the results reported by Lartey and Oferi (2000) and Egbe *et al*., (2010) in Ghana and Nigeria respectively for different cowpea accession, there also reported that the variation may be due to genotypic and environmental interactions.

In general, high level of genetic variability was observed in traits among the cowpea accession. According to Memo *et al*., (2005), the expression of the traits depends on the genetic characteristics of the planting materials under consideration as well as the environment in which they are grown. The non-significance of some of the traits may be due to less contributions to their development or might be due to genetic differences in the breeding materials evaluated. The significant association of some traits are expected for example, total pod weight was positively correlated with total seed weight of cowpea, this trait is an important parameter that determines the seed yield. This positive and significant association of these traits will provide plant breeders an understanding of phenotypic traits and their degree of association to be able to plan breeding schemes and managements of plant germplasm. Selection of highly associated traits such as total pod weight is an important trait than can be improved in the cowpea improvement program. Knowledge of the relationship among plant characters is useful while selecting traits for yield improvement. The negative correlation of yield with hundred seed weight is in disagreement with previous reports (Arshad *et al*., 2006; Basavaraja *et al*., 2005; Jyoti and Tyagi 2005; Banger *et al*., 2003) who all reported positive correlation between hundred seed weight and total Seed weight. From the result it can be inferred that direct selection for seed yield could be done for traits that are positively correlated to yield such as total pod weight, while indirect selection should be done for negative correlated traits like hundred seed weight. Path analysis revealed that seed yield can be improved by practicing selection for total pod weight, days to ninety-five percent maturity and numbers of braches per plant as they contribute directly to seed yield as revealed from the path analysis. It indicates the possibilities of simultaneous improvement of these traits by selection. This in turn will improve the seed yield, since they are positively correlated with seed yield. This present finding is also in similar trend of results reported by Singh *et al*., (1990), Kutty *et al*., (2003), Diriba, Shanko *et al*., (2014) and Mahesh *et al*., (2017) who reported direct and positive effects numbers of pods per plant, harvest index and biological yield on seed yield.

## CONCLUSIONS

The findings of this study shows that significant positive correlation exist between “branches per plant and pods per plant”, “pods per plant and Pod weight” as well as between “pod weight and seed weight”. Path coefficient analysis shows that pod weight has the highest direct contribution to yield while number of pods contributes indirectly to yield through pod weight. The study also identified genotype IT99K-573-1-1 as the best performing variety with yield of 930 kg ha-^1^. Based on the information obtained from this research, breeders can increase yield in cowpea by increasing pod weight. Further studies and multi-locational trials are recommended to consolidate this work, to enhance adoption by farmers.

## ACKNOWLEDGMENTS

We extend our heartfelt appreciation to the staff of the Molecular Biology Laboratory at Joseph Sarwuan Tarka University, Makurdi, Benue State. Additionally, our gratitude goes to the dedicated team at the Genetic Resource Center, International Institute of Tropical Agriculture (IITA), located in Ibadan, Nigeria.

